# Variations in nonlocal interaction range lead to emergent chase-and-run in heterogeneous populations

**DOI:** 10.1101/2024.06.17.599461

**Authors:** K. J. Painter, V. Giunta, J. R. Potts, S. Bernardi

## Abstract

In a chase-and-run dynamic, the interaction between two individuals is such that one moves towards the other (the chaser), while the other moves away (the runner). Examples can be found in both interacting cells and interacting animals. Here we investigate the behaviours that can emerge at a population level, for a heterogeneous group that contains subpopulations of chasers and runners. We show that a wide variety of patterns can form, from stationary patterns to oscillatory and population-level chase-and-run, where the latter describes a synchronised collective movement of the two populations. We investigate the conditions under which different behaviours arise, specifically focusing on the interaction ranges: the distances over which cells or organisms can sense one another’s presence. We find that when the interaction range of the chaser is sufficiently larger than that of the runner – or when the interaction range of the chase is sufficiently larger than that of the run – population-level chase-and-run emerges in a robust manner. We discuss the results in the context of phenomena observed in cellular and ecological systems, with particular attention to the dynamics observed experimentally within populations of neural crest and placode cells.

## 1 Introduction

In cellular and animal systems, interactions frequently trigger movement responses. The collective movements that emerge at a population level have become the focus of considerable interest, in phenomena that range from the swarming and flocking of animals [1], to embryonic development [2], and cancer invasion [3]. Mathematical modelling has helped uncover the mechanistic basis of these emerging dynamics, using methods that range from agent-based (interacting particle systems) to continuous PDE systems [4, 5, 6].

Attraction and repulsion form two fundamental interaction types, whereby the nearby presence of another individual induces movement towards (attraction) or away from it (repulsion). Within a single homogeneous population, attracting interactions can drive a population to self-organise into an aggregated group, such as a herd of animals, while repelling interactions can enhance the dispersal of an aggregated population. For multiple or heterogeneous populations – a mixture of distinct animal species or cell types, or subpopulations with different traits – greater complexity is possible [7]. In a binary system with two distinct populations, there are four principal interactions: a set of two homotypic interactions between individuals of same type, and a set of two heterotypic interactions between individuals of different type. A broad set of interaction combinations can be conjured and the question as to how these subsequently translate into population level dynamics are of manifest interest when it comes to understanding how heterogeneous populations become spatially structured. Beyond the fundamental nature of an interaction – whether it is attracting or repelling – a second important point of consideration is its range: the distance of separation over which an interaction can occur. These distances will naturally depend on the mechanism through which an interaction is mediated. Animals and cells can use a multitude of mechanisms to sense other individuals [8], through both direct and indirect means. By direct we mean a sense that could (almost) exactly signal the current position of another individual, for example through directly touching or sighting the neighbour; indirect refers to detection through an intermediary, such as chemical cues or tracks left by the neighbour, that may indicate its recent presence. Whether a sensing is direct or indirect, within heterogeneous groups the interaction ranges of the homotypic and heterotypic interactions could vary considerably: animals range widely with respect to the form and range of their sensory systems – certain species of baleen whales are believed to be possible of communicating over 100s of kilometres [9]; different cells can extend a spectrum of cell protrusions from shorter to long range [10].

We will focus here on a particular type of heterotypic interaction, which we call *chase-and-run* (or pursue-and-escape). Here, individuals of (say) the first population move towards individuals of the second population, which in turn move away. While for the most part we eschew a specific application in favour of broader insights, we note that such dynamics feature within numerous contexts. In an ecological context the obvious example would be between a predator and its prey [11], but it could also occur between dominant and submissive members of a pack. At the cellular level, chase-and-run has been observed both in vitro and in vivo for a variety of heterogeneous cell groups, such as between distinct zebrafish pigment cell types [12] or between embryonic neural crest and placode populations [13].

The case of neural crest and placode cell interactions offers a particularly illuminating exemplar in the context of the objectives here. Chase-and-run in this system is manifested at the population level, demonstrated through in vitro experiments in which a smallish (circa 100 cells) cluster of neural crest cells are cultured close to a similarly-sized cluster of placode cells [13]. These two clusters are subsequently observed to move in concert, maintaining a similar distance of separation as the neural crest cell cluster persistently pursues the cluster of placode cells. Given that certain pathologies may arise from failed migration of neural crest cells to a target tissue [14], the robustness of collective migrations of this nature may be crucial for correct tissue development.

Logically, population-level chase-and-run would seem a natural outcome of chase-and-run interactions at the individual level. However, it would seem equally plausible to suppose other possible outcomes: for example, a running cluster may escape a chasing group and move out of range, or chasers may catch the escaping group. Alternatively, individuals could be chasing one another in a range of different directions, causing the overall populations to be more-or-less homogeneously spread. The objective here is to systematically explore the various dynamics that can emerge within systems where a chase-and-run interaction occurs. To achieve this, in the next section we will describe a somewhat minimal population-level model whereby each interaction is characterised by its *type* (attracting or repelling), *strength*, and *range*. We show that this simple model is capable of exhibiting a broad spectrum of population-level dynamics, including those described above. We subsequently explore the conditions under which particular types of pattern can robustly emerge, focussing on the crucial role played by the interaction ranges.

## 2 Methods

We present our model as a system of nonlocal advection-diffusion PDEs for continuous population densities, a structure commonly used to describe interaction-based movements in cellular and animal populations (see [6] for a review). However, it is noted that these equations can be derived through coarse-graining a stochastic individual (agent-based) model, see Supplementary Information (SI); this connection is exploited later to determine the extent to which behaviours observed at the continuous (deterministic) level translate to the discrete (stochastic) level.

The model assumes that movements are governed by a set of homotypic and heterotypic interactions between neighbouring members of two populations: *chasers, C*, and *runners, R*. Neighbouring here refers to those members within a distance specific to a particular interaction. Precisely, we consider (see Figure 1(a)):

**Figure 1:**
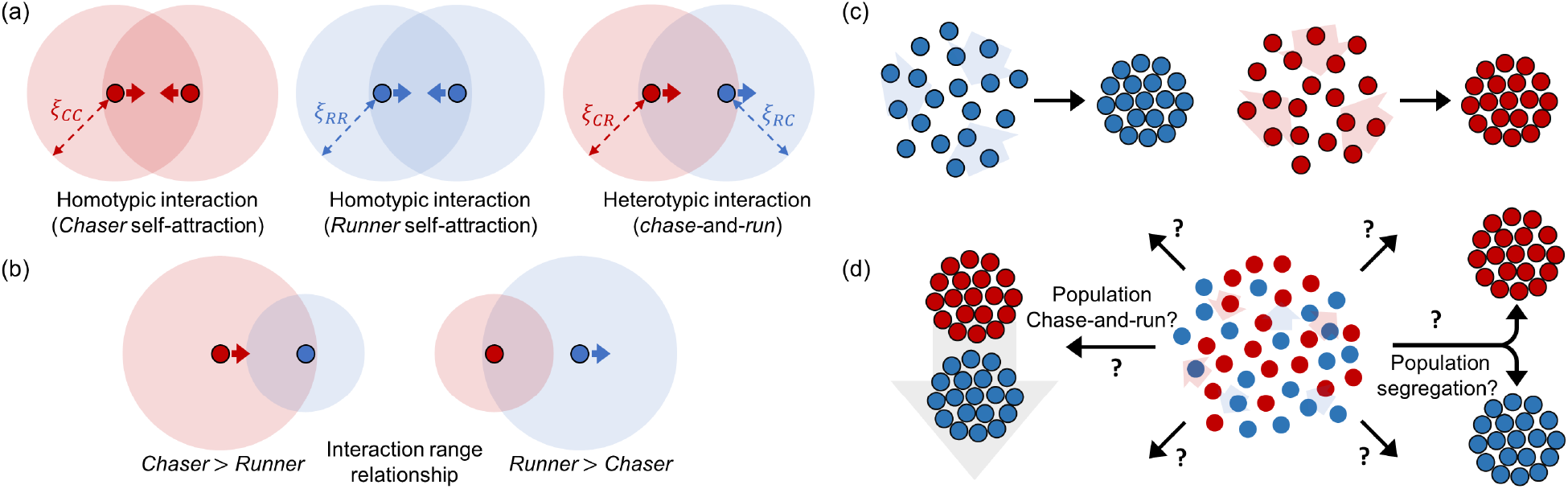
(a) Individual *chasers* (dark red circles) and *runners* (dark blue circles) interact across range *ξ*_*ij*_ (light coloured circles) such that (left) chasers are attracted to other chasers, (middle) runners are attracted to other runners, and (right) chase-and-run describes the heterotypic interactions. (b) Distinct ranges lead to various possibilities, such as distances over which (left) chasers detect runners but not vice versa, or (right) runners detect chasers but not vice versa. (c) In the absence of the other population, self-attracting interactions aggregate populations. (d) The principal question we ask in this paper is as follows: in a mixed scenario, what dynamics can arise and what drives the selection of a particular dynamic? Logical reasoning would suggest possibilities that could include a segregated scenario, where the two populations separate into two essentially non-interacting groups, or a population-level chase-and-run dynamic.

- Attracting homotypic interactions. This could represent animal species with naturally tendencies to herd or pack with conspecifics, or cell populations that self-adhere.
- Chase-and-run heterotypic interactions. Specifically, members of *C* are attracted to neighbouring members of *R* (the chase), but members of *R* are repelled by neighbouring members of *C* (the run).

Denoting by *C*(*x, t*) and *R*(*x, t*) the two population densities at position **x** *∈* Ω *⊂* ℝ^*n*^ and time *t ∈* [0, *∞*), we consider the following system of equations:

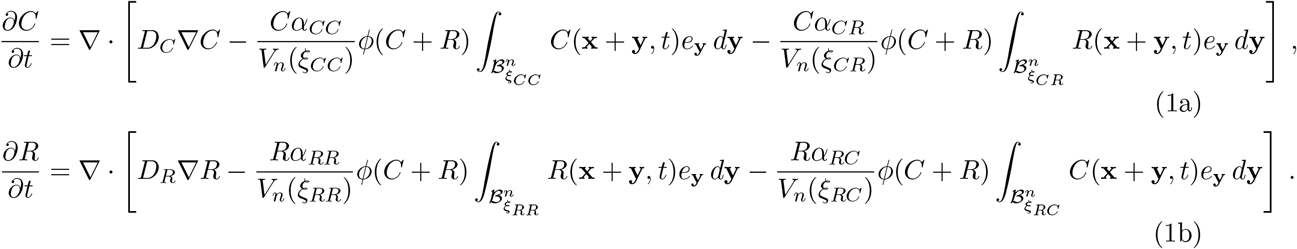

The three movement terms on the right hand side of (1a) correspond to an undirected (diffusive) component, a directed movement due to homotypic interactions between members of *C*, and a directed movement due to heterotypic interactions. Correspondingly, those on the right hand side of (1b) derive from diffusive movement, homotypic interactions between members of *R*, and heterotypic interactions. We use 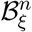 to denote the *n*-dimensional ball of radius *ξ, V*_*n*_(*ξ*) to denote its corresponding volume, and set *e*_**y**_ as the unit vector in the direction of **y** (if *n* = 1, this is understood to be the sign of *y*). Models with the above structure have been readily adopted in recent years – see the review [6] and we note in particular a number of studies for two (or more) species for both cellular [15, 16, 17, 18, 19, 20, 21] and ecological [22, 23, 24, 25, 26] interactions.

Parameters *α*_*ij*_, for *ij ∈ {CC, CR, RC, RR}*, measure the strength and form of response; specifically, the response of members of population *i* to members of populations *j*. A positive (negative) value for *α*_*ij*_ indicates that *i* is attracted to (repelled by) *j* and the magnitude determines the strength of response. Accordingly, we concentrate on a regime in which *α*_*CC*_, *α*_*RR*_, *α*_*CR*_ *≥* 0, but *α*_*RC*_ ≤ 0. The parameters *ξ*_*ij*_, for *ij ∈ {CC, CR, RC, RR}*, denote *interaction range* parameters: the maximum separation distance over which each type of interaction can occur. Therefore, a relationship *ξ*_*CR*_ *> ξ*_*RC*_ implies that the range over which the chase occurs is greater than that of the run (see Figure 1(b)). Note that we will often restrict to a two-dimensional parameter space (*ξ*_*R*_, *ξ*_*C*_), where *ξ*_*RC*_ = *ξ*_*RR*_ ≡ *ξ*_*R*_ represents the runner interaction range and *ξ*_*CR*_ = *ξ*_*CC*_ ≡ *ξ*_*C*_ represents the chaser interaction range. Restricting to these two parameters could be broadly interpreted as defining distinct perception ranges for each population: *ξ*_*R*_ (or *ξ*_*C*_) represents the perception range of runners (or chasers), and subsequently defines the maximum range over which its homotypic and heterotypic interactions can form. As a note, for simplicity the model relies on a “top hat” interaction kernel (for other kernels, see [6] and references therein), in which the magnitude of an interaction response does not change with the separation distance (beyond individuals needing to be within the relevant range).

We note that *D*_*C*_ and *D*_*R*_ are diffusion coefficients that measure the level of random movement. The function *ϕ*(*C* + *R*) curbs directed movement if the population density becomes excessively high. This is not an issue in one dimension, where we will therefore set *ϕ*(*C* + *R*) = 1 for simplicity. In two dimensions, however, self-attraction tends to generate highly concentrated population densities when *ϕ*(*C* + *R*) = 1, and we will set *ϕ*(*C* + *R*) = 1*/*(1 + *C* + *R*) to avoid this unwanted behaviour in 2D.

We study (1) in both 1D and 2D geometries. To minimise domain/boundary-induced effects we impose wrap-around (periodic) boundary conditions, by setting the domain Ω to be a circle (of circumference *L*) in 1D, or torus (of dimensions *L× L*) in 2D. Under these boundary conditions the total population masses are conserved and uniform steady state distributions are therefore determined by the mean initial distributions, i.e. 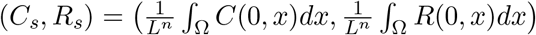. Two forms of initial conditions are considered: *pre-aggregated*, where *C*(0, *x*) and *R*(0, *x*) are concentrated as Gaussian distributions; *dispersed*, where *C*(0, *x*) and *R*(0, *x*) are uniform, bar a small random perturbation. For convenience we assume identical diffusion coefficients and mean initial densities and an *a priori* scaling of length, time and densities such that *C*_*s*_ = *R*_*s*_ = *D*_*C*_ = *D*_*R*_ = 1.

## 3 Results

### Chase-and-run generates a broad spectrum of population dynamics

To motivate the investigation that follows, we pose a simple question: does chase-and-run at an individual level lead to chase-and-run at the population level? More precisely, can aggregated groups of the two populations move in synchronicity (see Figure 1(d))? Logically, positive homotypic interactions should allow runners and chasers to maintain aggregated forms, and the heterotypic chase-and-run should coordinate the group movements. Simulations confirm this natural conceit, where we observe that two groups placed in proximity (such that a non-negligible level of heterotypic interactions is initially present) form a sustained synchronised movement, see Figure 2(a).

**Figure 2:**
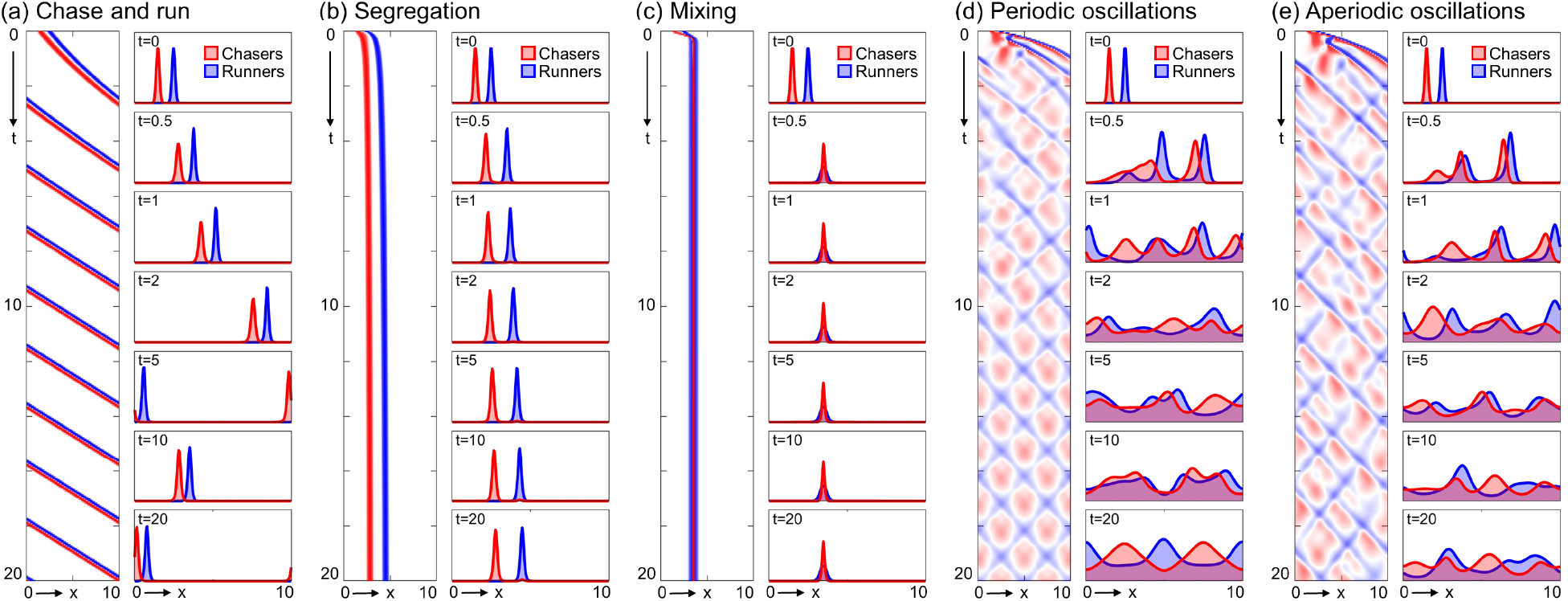
Dynamics of the chase-and-run system (1) in 1D on [0, 10]: each panel shows chasers (red) and runners (blue) on a kymograph and distributions at the stated times. Here interaction ranges *ξ*_*RR*_ = *ξ*_*CR*_ = *ξ*_*RC*_ = *ξ*_*RR*_ = 1 and interactions strengths *α*_*CC*_ = *α*_*RR*_ = 3 with: (a) (*α*_*CR*_, *α*_*RC*_) = (1, −1), (b) (*α*_*CR*_, *α*_*RC*_) = (2, −3), (c) (*α*_*CR*_, *α*_*RC*_) = (2, −1), (d) (*α*_*CR*_, *α*_*RC*_) = (5, −5), (e) (*α*_*CR*_, *α*_*RC*_) = (4, −4). Note that *D*_*C*_ = *D*_*R*_ = 1 with initial conditions *C*(0, *x*) = *C*^∗^ exp (−50(*x* − 1.5)^2^), *R*(0, *x*) = *R*^∗^ exp (−50(*x* − 2.5)^2^), where *C*^∗^ and *R*^∗^ are set such that *C*_*s*_ = *R*_*s*_ = 1.

However, this (numerically) stable configuration is found to break down under relatively subtle variations to the interaction strengths. For other parameter combinations we observe population segregation. Here, runners escape the interaction range of chasers and the two populations settle into fixed and separated aggregates, see Figure 2(b), each supported through self-attraction. An alternative form of stationary pattern occurs when chasers are able to reach and trap the runners, resulting in a single mixed aggregate, see Figure 2(c). More complicated dynamics arise for other parameter combinations where, rather than settling into stable chase-and-run or stationary aggregates, we observe sustained oscillations in space and time, which can be both periodic (Figure 2(d)) or aperiodic (Figure 2(e)). Having established that a broad spectrum of patterns are possible, we next explore how interaction range impacts on emergent dynamics in populations of chasers and runners.

### Interaction ranges determine pattern selection

As noted, different interactions may operate across different ranges, e.g. if populations have distinct limits to their perception depth. We consider emergent dynamics across (*ξ*_*R*_,*ξ*_*C*_) parameter space from dispersed initial conditions, which allows us to determine whether a particular pattern forms without initial bias. Then, a Turing-type linear stability analysis (LSA) can be used to establish whether the uniform steady state can be unstable to inhomogeneous perturbations and patterns emerge (see SI). Largely speaking, homotypic attractions drive pattern formation – although we note some subtleties that emerge in two dimensions [20] – and allow one or both of the populations to accumulate into one or more groups. The presence of a heterotypic chase-and-run typically leads to a ‘Turing-wave’ type instability, so that emerging patterns are predicted to oscillate in both space and time; these oscillations are observable in the early time dynamics for each of the simulations plotted in Figure 3(a)ii-v.

**Figure 3:**
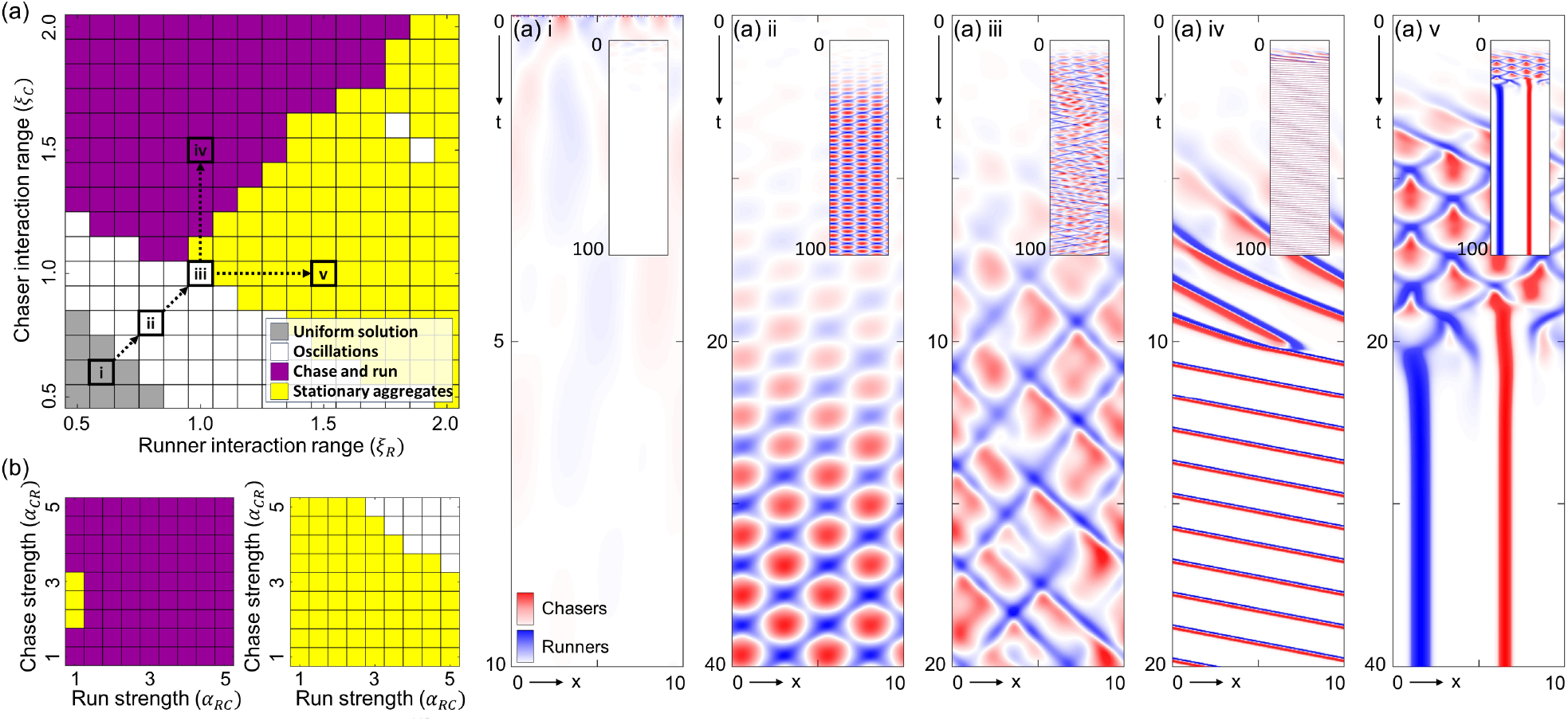
(a) Pattern selection across interaction range space. We classify the pattern for each (*ξ*_*R*_, *ξ*_*C*_) pair as: uniform solution (gray), spatiotemporal oscillations (white), population chase-and-run (magenta), stationary aggregates (yellow). The kymographs plotted in (i-v) illustrate representative solutions for certain (*ξ*_*R*_, *ξ*_*C*_) pairs, as indicated in (a). For other model parameters, we fix *α*_*CC*_ = *α*_*CR*_ = −*α*_*RC*_ = *α*_*RR*_ = 3, *D*_*C*_ = *D*_*R*_ = 1, *L* = 10 and initial conditions in the dispersed form, such that *C*_*s*_ = *R*_*s*_ = 1. (b) Pattern selection across interaction strength space. Here, the pair (*α*_*RC*_, *α*_*CR*_) is varied, whilst interaction ranges are selected (left) from the chase and run region (specifically, (*ξ*_*R*_, *ξ*_*C*_) = (1, 1.5)), and (right) from the stationary aggregates region (specifically, (*ξ*_*R*_, *ξ*_*C*_) = (1.5, 1)). We fix *D*_*C*_ = *D*_*R*_ = 1, *L* = 10 and initial conditions in the dispersed form such that *C*_*s*_ = *R*_*s*_ = 1.

LSA provides insight into initial pattern emergence, but for longer time dynamics we rely on numerical simulations. We numerically solve (1) for different (*ξ*_*R*_,*ξ*_*C*_) combinations and classify the dynamics at the end of each simulation. We cannot exclude that an observed pattern represents a long-time transient rather than a stable form [27], but simulations are performed for times that extend significantly beyond the characteristic timescale of pattern formation (an order of magnitude or more). Figure 3(a) shows the classification of parameter space, where we observe a clear demarcation into distinct regions. If both species have small interaction ranges, patterns do not form and densities evolve to the uniform steady state solution, Figure 3(a)i – this region coincides with the region of stability predicted by LSA. Intuitively, small interaction ranges limit the size of the observable population and restrict how much interaction-based movement can be generated. Adjacent to this region we observe sustained spatiotemporal oscillations. These transform from periodic and low amplitude (Figure 3(a)ii) to aperiodic and large amplitude (Figure 3(a)iii) with increasing distance from the stability/instability boundary; this coincides with increased pattern growth rates and an increase in the number of unstable modes, predicted by the LSA (see SI).

Setting even larger interaction ranges results in patterns that initially oscillate before transitioning into one of two general forms: a population-level chase-and-run (Figure 3(a)iv) or stationary aggregates (Figure 3(a)v). Significantly, the regions where these two pattern types develop depend on the interaction range relationship. Broadly speaking, population chase-and-run emerges when the chasers have a (sufficiently) larger interaction range than the runners, whereas stationary aggregates form when runners have the larger interaction range. This division is more strongly accentuated when higher interaction strengths are used (see Figure S2(a)), where we find chase-and-run (*ξ*_*C*_ *> ξ*_*R*_) or stationary aggregates (*ξ*_*C*_ *< ξ*_*R*_) and greatly reduced regions of oscillating or uniform solutions. To test whether it is the heterotypic interactions that drive this pattern selection, we see a similar division when only (*ξ*_*CR*_, *ξ*_*RC*_) are varied: *ξ*_*CR*_ *> ξ*_*RC*_ (*ξ*_*RC*_ *> ξ*_*CR*_) biases pattern selection to a chase-and-run (stationary) form (Figure S2(b)).

To probe further how the nature of the chase-and-run interactions drive the dynamics, we investigate pattern selection in each of two regimes *ξ*_*C*_ *> ξ*_*R*_ and *ξ*_*R*_ *> ξ*_*C*_, while changing (*α*_*RC*_, *α*_*CR*_). This allows a test into the robustness of an observed dynamic while modulating the strength of the heterotypic interactions. Corroborating the key importance of interaction range, we see a robust emergence of chase-and-run across a wide range of interaction strengths, if *ξ*_*C*_ *> ξ*_*R*_ (Figure 3(b), left panel). Similarly, we see robust emergence of stationary aggregates when *ξ*_*R*_ *> ξ*_*C*_ (Figure 3(b), right panel). These explorations also highlight some further subtleties. For example, stationary aggregates are found to range from completely segregated (clusters of only chasers or only runners), to mixed/segregated (some clusters contain both chasers and runners), to completely mixed (the two populations are co-localised), see Figure S2(c). This transition follows changing strengths of the heterotypic interactions: *α*_*RC*_ *> α*_*CR*_ can allow runners to completely escape chasers and form a separate group, while *α*_*CR*_ *> α*_*RC*_ can allow chasers to catch and trap runners. Modulations to the chase-and-run can also occur. For example, lower run interaction strength results in some runners being left behind; on the slightly artificial periodic domain, these runners are subsequently recollected and the process repeats (Figure S2(d)).

Summarising, we find that distinct interaction ranges have a strong influence on emergent population level dynamics under chase-and-run type interactions.

### Chase-and-run in the plane

In two dimensions a runner can escape a chaser across all angles of the circle. We therefore investigate how dynamics change for populations distributed across Ω = [0, *L*] *×* [0, *L*], under the same general homotypic and heterotypic interactions. Populations are initially dispersed (Figure 4(b)) and we again explore (*ξ*_*R*_, *ξ*_*C*_)-space. Simulations in this section are accompanied by SI animations.

**Figure 4:**
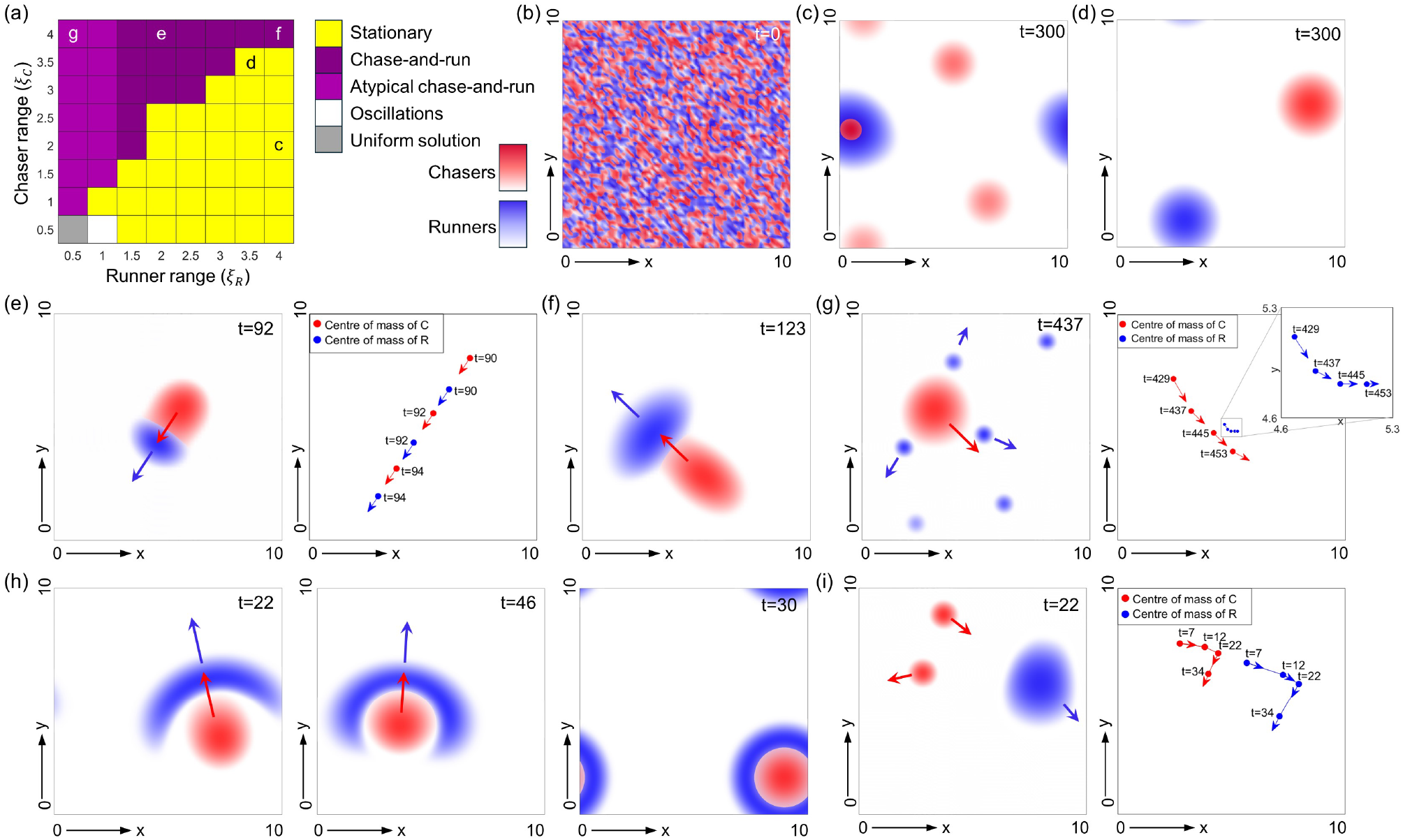
(a) Classification of (*ξ*_*R*_, *ξ*_*C*_) parameter space, for simulations of (1) in 2D. (b) Dispersed initial conditions at *t* = 0: random perturbations about (*C*_*s*_, *R*_*s*_) = (1, 1). (c) Stationary aggregate solution found at point **c** in (a). (d) Stationary aggregate solution found at point **d** in (a). (e, left) Stereotypical chase-and-run found at **e** in (a), see Supplementary Movie 1; (e, right panel) The positions of the centre of masses for chasing and running groups over a range of times. (f) Chase-and-run, with interaction ranges found at **f**, see Supplementary Movie 2. (g, left panel) Atypical chase-and-run found at **g** in (a), see Supplementary Movie 3; (g, right panel) Centre of masses for the chasing and the running groups. (h) Crescent-shaped runners for *ξ*_*CC*_ = *ξ*_*CR*_ = *ξ*_*RR*_ = 4 and (left) *ξ*_*RC*_ = 2, (middle) *ξ*_*RC*_ = 1, (right) *ξ*_*RC*_ = 0.5. See Supplementary Movies 4-6. (i, left panel) Atypical chase-and-run for *ξ*_*CR*_ = *ξ*_*RC*_ = *ξ*_*RR*_ = 4 and *ξ*_*CC*_ = 1, see Supplementary Movie 7; (i, right panel) Centre of masses for the chasing and the running groups. All simulations in this figure use *α*_*CC*_ = *α*_*RR*_ = *α*_*CR*_ = −*α*_*RC*_ = 15, *D*_*C*_ = *D*_*R*_ = 1, *L*_*x*_ = *L*_*y*_ = 10 and *ϕ*(*C* + *R*) = 1*/*(1 + *C* + *R*).

Principally, we observe the same separation of parameter space into sustained population-level chase-and-run (when *ξ*_*C*_ *> ξ*_*R*_) or stationary aggregates (when *ξ*_*R*_ *> ξ*_*C*_), see Figure 4(a). Consistent with 1D, a more defined separation emerges with higher interaction strengths: lower interaction strengths lead to smaller regions of chase-and-run or stationary aggregates (see Figure S3(a, top)), and larger regions of oscillating (see Figure S3(b)i for an example) or uniform solutions. While chase-and-run is implemented through the heterotypic interactions, increasing the chase-and-run strength alone does not encourage emergence of these dynamics at the population level; in contrast, we see an expansion of the oscillating regime (see Figure S3(a, bottom)). This reinforces that both homotypic and heterotypic interactions play a critical role in setting the population level dynamics.

Consistent with 1D, we find that stationary aggregates range from mixed (Figure 4(c)) to segregated (Figure 4(d)). In regions of chase-and-run, homotypic self-attraction first organises the dispersed populations into clusters. Then, heterotypic interactions drive the cluster of chasers to pursue the cluster of runners, maintaining consistent shape and speed (Figure 4(e), Supplementary Movie 1). This stereotypical chase-and-run robustly emerges when self-interaction strengths are sufficiently strong, and on larger domains (Figure S3(c)). However, some subtler features become apparent according to the runner interaction range. Under more-or-less equal interaction ranges the two groups are roughly equal in size (Figure 4(f), Supplementary Movie 2). Reducing the runner interaction range groups runners into a smaller (and more concentrated) group (Figure 4(e)). Below some critical value, however, runners instead organise into multiple small groups. Run-and-chase still occurs, but in an atypical form: chasers continue to pursue runners, but with multiple runner clusters the chasing direction changes over time (Figure 4(g), Supplementary Movie 3). With its limited interaction range, a cluster of runners remains fixed in position until a chasing group is sufficiently close, at which point it moves away.

To investigate further patterning subtleties, we perturbed specific interaction range parameters and comment on notable instances. Lowering only the run interaction range (*ξ*_*RC*_) leads to a crescent-shaped configuration ahead of the chasing group (Figure 4(h), Supplementary Movies 4-5). Here, runners only detect the nearest portion of the chasing group and, consequently, the escape direction can vary significantly across the runner population. Self-attraction of runners (*ξ*_*RR*_) occurs over longer range, maintaining cohesion and generating the crescent. When *ξ*_*RC*_ decreases below a critical value, runners entirely surround chasers in an annulus-like shape, so that net motion is prevented (Figure 4(h), Supplementary Movie 6). Finally, reducing the chaser self-attraction range (*ξ*_*CC*_) below a critical value results in multiple (smaller) chaser groups. At a phenomenological level, the subsequent dynamics can be viewed as a reversal of the atypical chase-and-run above, where now a single group of runners is chased by multiple groups of chasers (Figure 4(i), Supplementary Movie 7). These results highlight that the individual homotypic interaction ranges strongly influence the spatial extension of groups at the population level.

Broadly speaking, investigations in 2D reinforce that interaction ranges dictate the emerging dynamics. Yet we also observe increasingly nuanced patterning, suggesting that greater control is required if a particular pattern is to emerge in higher dimensions.

### Individual-based modelling

The continuous model is predicated on an assumption that the populations can be approximated by density distributions, rather than individual positions. The extent to which this is valid for the population sizes encountered in real-world instances of collective movement is uncertain, and we now determine whether behaviours observed for (1) carry to an individual-based model (IBM). The lattice-based IBM is described in the SI, and features two populations of particles which interact according to the homotypic and heterotypic interactions described previously. Notably, coarse-graining the IBM leads to the continuous formulation given by Equation (1) (see SI). Consequently, an IBM parameter set (including lattice spacing and time step) can be chosen in correspondence to a given simulation of the continuous model. Note that for all IBM simulations we use 100 individuals per populations (consistent with typical sizes of populations of neural crest and placode cells in the example cited in the introduction).

For the 1D IBM (see SI) we consider a lattice of length *L* = 10 with spacing *l* = 0.1. Setting the time step *τ* = 0.005 then ensures *D* = 1, as used in simulations of the continuous models. As before, we keep *α*_*RC*_ *<* 0 and *α*_*CC*_, *α*_*CR*_, *α*_*RR*_ *>* 0, so that individuals of population *C* chase those of population *R*. In Figure 5(a) we classify the type of pattern observed following simulations of the IBM, covering the same parameter space as in Figure 3(a). Consistent with the continuous model, we see sustained chase-and-run when the interaction range of chasers is large compared to those of the runners (Figure 5(b)), otherwise we see more-or-less stationary aggregates (Figure 5(c)); there can be some ‘wobble’ in a stationary aggregate position over time, due to inherent stochasticity, but there is no clear chasing. Finer points of detail also translate, such as the range of stationary aggregate patterns from completely segregated to mixed. We further confirmed through simulations of a 2D IBM, using equivalent settings to simulations of the continuous models reported in Figure 4. A population chase-and-run behaviour can be observed when *ξ*_*C*_ *> ξ*_*R*_ (Figure 5 (d)), and stationary aggregates for *ξ*_*R*_ *> ξ*_*C*_ (Figure 5 (e)).

**Figure 5:**
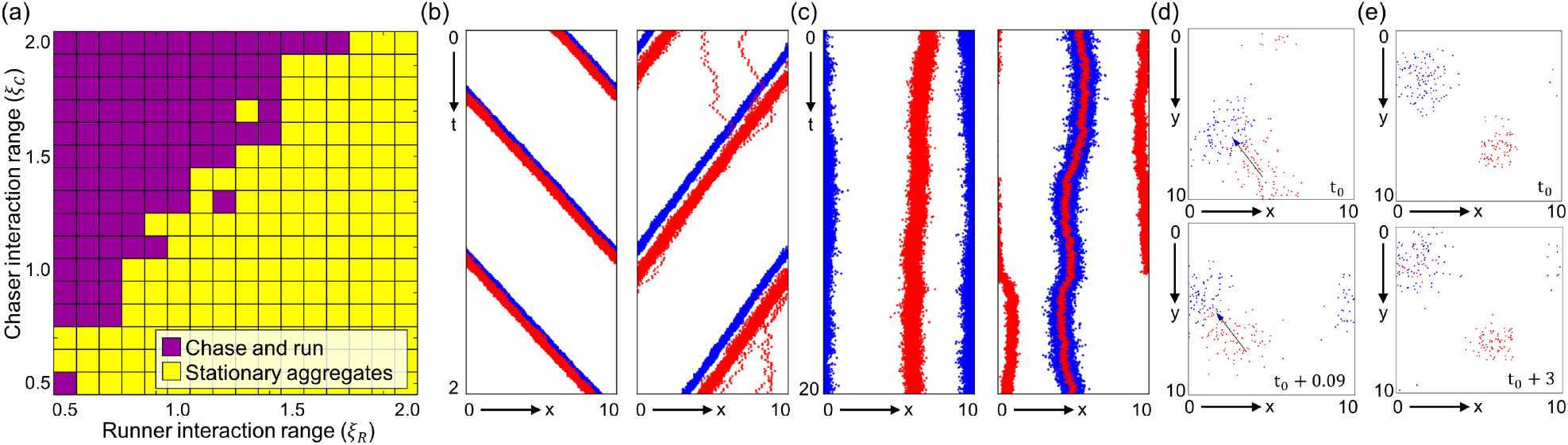
Simulation output of the IBM. (a)-(c) 1D simulations. (a) Classification of (*ξ*_*R*_, *ξ*_*C*_) space analogous to Figure 3(a). (b) Representative examples of chase-and-run dynamics in the IBM. (c) Representative examples of stationary aggregate dynamics in the IBM. (d-e) 2D simulations. (d) 2D chase-and-run, analogous to Figure 4(e) with plots showing chasers (red) and runners (blue) at a reference time (top) *t*_0_ and (bottom) *t*_0_ + 0.09. (e) 2D stationary aggregates analogous to Figure 4(c) with plots at a reference time (top) *t*_0_ and (bottom) *t*_0_ + 3.

However, some noteworthy differences emerge. First, for the IBM we have not observed (clear) patterns representing homogeneity or sustained oscillations, at least not when using parameter settings equivalent to those within the continuous simulations. This perhaps suggests that these patterns are more reliant on the assumption that a population can be approximated by a continuous distribution, rather than individual positions. Also, in 2D simulations we have found chase-and-runs where the direction of group movements change over time: at lower population sizes, the occasional loss of individuals from runner or chaser groups can prove influential, creating ‘rogue’ individuals that can subsequently alter cluster dynamics.

### Self-attraction is required for robust chase-and-run

We have have observed how population chase-and-run emerges in 1D, 2D and in IBM simulations, when the range of the chase is greater than that of the run. However, all simulations to date assume that each population has a self-attraction that allow their self-aggregation. We investigate whether this is crucial for the emergence and maintenance of a pattern type.

First we repeat the 1D analysis, investigating pattern selection in (*ξ*_*R*_, *ξ*_*C*_)-space under two cases: (i) only runners self-aggregate (*α*_*RR*_ *>* 0, *α*_*CC*_ = 0); (ii) only chasers self-aggregate (*α*_*CC*_ *>* 0, *α*_*RR*_ = 0). In both instances we (broadly) find the same behaviour: across a large region of parameter space corresponding to *ξ*_*C*_ *> ξ*_*R*_, a sustained chase-and-run emerges at the population scale (Figure 6(a-b)). Self-aggregation in the runners leads to their clustering which – via the heterotypic interactions – collect chasers into a looser group in the rear (Figure 6(c), top). Self-aggregation in the chasers lead to a chaser cluster that ‘herds’ runners ahead (Figure 6(c), bottom).

**Figure 6:**
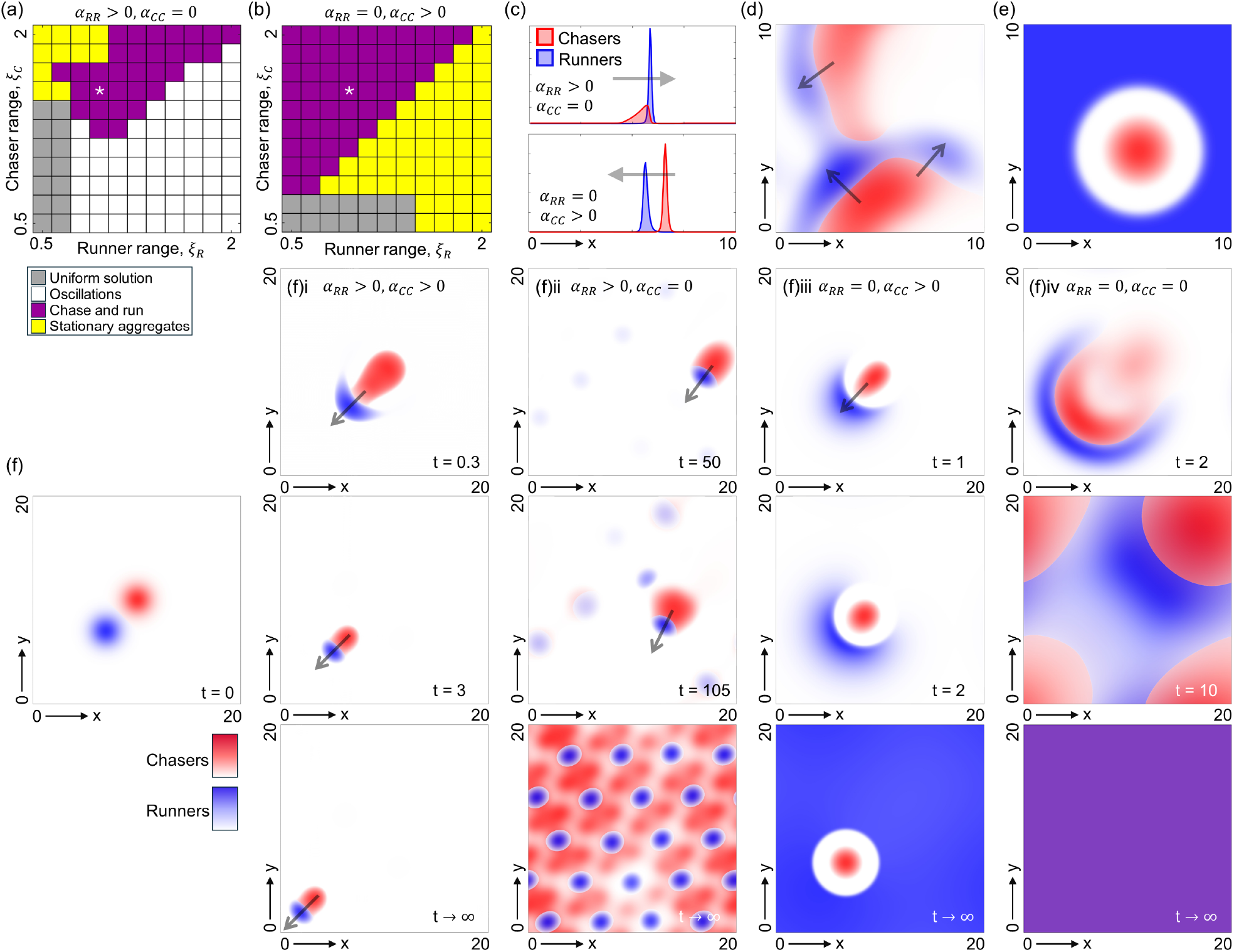
Dynamics for various self-aggregating tendencies. (a-b) Classification of (*ξ*_*R*_, *ξ*_*C*_)-space for (1) in 1D with *α*_*CR*_ = 3, *α*_*RC*_ = −3 and (*α*_*RR*_, *α*_*CC*_) = (a) (3, 0), (b) (0, 3). (c) Typical chase-and-run profiles in 1D, for positions marked with the white star in (a-b). (d) Irregular chase-and-run in 2D when only runners self-aggregate, from dispersed initial conditions (Supplementary Movie 8). Here, *ξ*_*C*_ = 4, *ξ*_*R*_ = 3, *α*_*RR*_ = *α*_*CR*_ = 15, *α*_*RC*_ = −15, *α*_*CC*_ = 0. (e) Stationary aggregate of chasers in 2D when only chasers self-aggregate. Here, *ξ*_*C*_ = 4, *ξ*_*R*_ = 2, *α*_*CC*_ = *α*_*CR*_ = 15, *α*_*RC*_ = −15, *α*_*RR*_ = 0. (f) Starting from pre-aggregated initial conditions (left panel), columns from left to right show snapshots for (*α*_*CC*_, *α*_*RR*_) = (i) (15, 15), (ii) (0, 15) (Supplementary Movie 9), (iii) (15, 0) (Supplementary Movie 10), (iv) (0, 0). Other parameters set at *ξ*_*C*_ = 4, *ξ*_*R*_ = 2, *α*_*CR*_ = 15, *α*_*RC*_ = −15 and “*t* → *∞*” indicates the long time pattern form. All simulations use *D*_*C*_ = *D*_*R*_ = *C*_*s*_ = *R*_*s*_ = 1.

However, these observations of emergent and sustained chase-and-run in 1D evaporate in higher dimensions. When only runners aggregate, scenarios of chase-and-run (*ξ*_*C*_ *> ξ*_*R*_) become ‘incoherent’, with the groups repeatedly splitting and reforming over time (Figure 6(d), see supplementary Movie 8). Here, the somewhat dispersed chasing group transmits variable directional information into the cluster of runners, inducing their escape into directions that can split the group; the reformation of groups can be part-attributed to the peculiarities of the boundary conditions. When only chasers aggregate, a *ξ*_*C*_ *> ξ*_*R*_ scenario leads to a stationary cluster of chasers surrounded by uniformly dispersed runners (Figure 6(e)). Unlike in 1D, where runners can be herded and trapped by the geometry, different escape directions mean that runners spread until chasers have no coherent direction of pursuit.

To test this under a more controlled setting, chasers and runners are initialised as juxtaposed clusters, Figure 6(f): this could be regarded as an optimal initial state to encourage chase-and-run. Also, to limit the influence of boundary conditions we consider a larger domain. Setting *ξ*_*C*_ *> ξ*_*R*_, if both populations self-aggregate we observe robust and coherent chase-and-run in (seeming) perpetuity (Figure 6(f)i). When only runners self-aggregate a transient chase-and-run can occur, but the runner and chaser groups lose mass over time and eventually chase-and-run disintegrates (Figure 6(f)ii, see Supplementary Movie 9). When only chasers self-aggregate, any chase-and-run is extremely brief: the runners quickly disperse in all directions to leave a stranded group of chasers (Figure 6(f)iii, see Supplementary Movie 10). Similarly, when neither population can self-aggregate any coherent motion is transient and the populations disperse to the uniform steady state (Figure 6(f)iv).

## 4 Discussion

We have explored the collective dynamics that emerge within mixed populations of chasers and runners, where the two populations have an inherent tendency to aggregate with members of their own kind and pursue and evade the other population, respectively. Focussing on the interaction ranges, we observe two dominating pattern forms: (i) when runners interact over larger distances than chasers (or the range of the run is greater than that of the chase), quasi-stationary and separated clusters emerge that undergo negligible interaction; (ii) when chasers interact over larger distances than runners (or the range of the chase is greater than that of the run), a collective chase-and-run forms where the cluster of chasers pursues the cluster of runners. Given sufficiently strong interaction strengths, this overall observation is apparently robust with respect to moderate parameter variations, when extended into a higher dimension setting, and when examined within the finite population sizes and stochastic setting of a corresponding individual-based model. As such, distinct interaction ranges between species appear to significantly determine emergent collective dynamics.

Intentionally, we have formulated the model in a minimal and generic manner, so that (in principal) the results could be interpreted for cellular or animal populations. For the former, chase-and-run dynamics have been observed in zebrafish pigment cells [12] and xenopus neural crest cell (NCC)–placode cell (PC) populations [13]. The latter forms a particularly apposite example, as *in vitro* studies reveal an analogous phenomenon of population-level chase-and-run [13]: NCCs form the chasing group, and PCs the running group. Experiments have uncovered various cellular interactions, including contact inhibition of locomotion (CIL) [28, 13] between all cells: the touching of two cells reverses their movement directions. Consequently, CIL forms a repulsion between all cells, but other interactions can switch some of these to attracting. First, PCs adhere to each other (through E-cadherin), which overcomes CIL to allow PC-to-PC attraction[13]. Second, attracting homotypic interactions also occur between NCCs [29, 30], seemingly autotaxis-mediated through a secreted attractant. Third, NCCs are attracted to PCs through the latter secreting a chemotaxis factor (Sdf-1) [13]. The net interactions may therefore conform to those explored here and we predict that a robust population chase-and-run will arise if NCCs interact over larger ranges. Indeed, this is quite plausible: PC interactions are seemingly through direct contact (membrane-membrane binding), while NCC interactions involve diffusing ligands. The simulations presented in Figure 6(f) are particularly apposite for this application, as the *in vitro* experiments place NCCs and PCs as juxtaposed clusters (akin to the *t* = 0 setting of those simulations). While indefinite chase-and-run requires both populations to have self-attraction, long transient chase-and-runs can still occur when only the runners have (strong) self-attraction: if this transient is much longer than the timescale of experiments, this may be sufficient. Thus, our simulations would suggest that while strong self-attraction in the runner group (i.e. the PCs) is essential, a similar requirement may not be strictly necessary for the chasers (i.e. the NCs). Fully investigating this system would be an interesting avenue, which could be achieved through extending to include the explicit interactions and key molecular factors; this would offer a complementary approach for modelling this system, as previous approaches have been restricted to agent-based models (e.g. [31]). In ecology, the model described here could help in understanding the spatial arrangements of predators and prey on a landscape. The concept of a ‘landscape of fear’, whereby prey avoid areas they believe predators to be living, has gained much attention in recent years [32, 33]. Improvements in animal tracking technology [34] have begun to allow researchers to map such landscapes and rigorously demonstrate prey avoidance [35, 36]. Likewise, the notion of prey-taxis [37], whereby predators seek to locate themselves near prey, is well-understood from both empirical [38] and mathematical [39] perspectives. Both aspects have been studied mathematically in a model similar to ours but with quadratic rather than linear diffusion [40]. However, in empirical situations, it is less clear whether a combination of predator-avoidance and prey-taxis behaviour ever actually leads to the kind of population-level chase-and-run described here. Indeed, empirical evidence suggests that most animals tend to locate themselves in relatively-stationary home ranges rather than moving over the landscape in response to predation pressure or seeking out mobile prey [41]. A possible reason for this is that landscape heterogeneity might ‘pin’ the population-level movements, so that, for example, prey might spend time in a woodland area where they can hide from predators, whereas predators might live close to the forest in hope of catching any prey that venture out [42]. Mathematical investigation of individual-level chase-and-run within heterogeneous environments would help understand why we still tend to see predators and prey using relatively stationary home ranges, despite the theoretical possibility of population-level chase-and-run in homogeneous environments.

Notions of followers and leaders have gathered considerable traction in recent years [43, 44, 45, 46, 3]. The terms ‘followers’ and ‘leaders’ imply a hierarchy – the latter guide the former – though in some cases their designations may simply reflect a spatial position [47]: leaders at the front, followers at the rear. Placed in the context of the collective run-and-chase that emerges here, the macroscopic perspective is runners at the front and chasers at the rear. But regarding runners as leaders and chasers as followers is clearly misleading: while the groups sort into a structured arrangement, there is no overall guidance and the net movement (from unbiased initial conditions) will be at random. This susceptibility is highlighted by the (2D) IBM simulations, where the chase-and-run direction was deflected by ‘rogue’ individuals and/or stochastic fluctuations. The addition of a further directional bias – such as a chemoattractant – could confer a robustness to the global direction of movement and in this regard it may be illuminating to observe how robustness alters according to which population detects the global cue: under what circumstances do runners or chasers make the more effective leaders?

Coherent chase-and-run – by which we mean a compact cluster of chasers pursuing a compact cluster of runners – was found to emerge in both 1D and 2D. In 1D this is particularly robust, given the stipulated relationship between interaction ranges, within both continuous and IBM formulations. Of course, this robustness can be partly attributed to the peculiarities of a 1D geometry, where runners can only escape in one of two possible directions. Away from singular or engineered instances – such as cells restricted to lines along microfabricated surfaces [48] – movements take place in higher dimensions, e.g. two dimensions for cells plated on a surface or animal movements on the plane. Despite the greater freedom movement direction, coherent chase-and-run does emerge robustly in two dimensions, but with caveats: for example (in continuous simulations) ‘complex’ chase-and-runs where the dynamics of a single large cluster of runners (chasers) is driven by multiple smaller groups of chasers (runners), or (in IBM simulations) with occasional fluctuations in the movement direction. Emergence of a particular dynamic in two dimensions may therefore require additional ‘control’, e.g. more tightly controlled parameters or reinforcement through environmental heterogeneity. A number of modelling and experimental studies have suggested that microenvironmental factors may indeed play an important role in confining collective migration of neural crest cells along particular pathways [49, 50].

We have observed even more diverse pattern forms in certain regimes, including a menagerie of oscillatory dynamics. These dynamics were robust – in the sense that they continued indefinitely and persisted under perturbation – and therefore seemingly represent a stable pattern outcome of the continuous model, rather than a transient dynamic. A deeper analytical study into the patterns that arise as parameters are varied could be achieved through nonlinear bifurcation analyses [51], a possible future direction. However, is is noted that in the IBM these oscillatory dynamics were conspicuous by their absence: under equivalent parameters, solutions evolved to either the population chase-and-run or stationary aggregate patterns. The continuous model assumes that populations can be approximated by a continuous density rather than individual particle positions, and it is possible that simulations of the IBM under larger populations would reveal similar phenomena. The IBM does of course introduce other forms of artificiality – for example, instantaneous jumps in individual location – and convergence with the continuous model involves various limits (e.g. smaller steps, higher population sizes): a more detailed investigation into the correspondence between the IBM and continuous model may pinpoint scenarios under which particular outcomes may be expected to arise. Overall, though, it reinforces the general lesson that caution should be adopted regarding ‘exotic’ patterning phenomena observed in models, particularly when found in restricted regions of parameter space.

A number of earlier studies have used similar modelling frameworks to understand the distinct patterns that can arise from self and cross interactions within heterogeneous systems. Several of these have been motivated by adhesive cell sorting, for example see [15, 52, 17, 18, 19, 21, 20]. Here, homotypic and heterotypic interactions are typically nonnegative and studies have focussed on how distinct adhesions lead to distinct arrangements, consistent with classical predictions of the differential adhesion hypothesis [53]. A generalisation to interactions that range from attracting to repelling was conducted in [16], although the subsequent analysis was primarily restricted to simple one-dimensional geometries and equivalent interaction ranges. Recently, Jewell et al [20] have extended this to two dimensions where, surprisingly, a pure chase-and-run interaction (with no homotypic interactions) was shown to be capable of generating pattern formation in two (and higher) dimensions. Significantly, unequal interaction ranges form a requirement and the study here further reinforces the critical role that interaction ranges can play in patterning dynamics.

We have shown that wide-ranging collective dynamics can emerge, even for just two species undergoing chase-and-run interactions. Yet, the relatively simple assumption of distinct interaction ranges provides a powerful point of control, strongly determining pattern selection. Clearer understanding into the ranges over which particular interactions are mediated would provide valuable information into the organisation of complex systems.

## Ethics

This work did not require ethical approval from a human subject or animal welfare committee.

## Data Access

Code for 1D simulations is available at https://github.com/kjpainter/ChaseAndRun. Code for 2D simulations is available at https://github.com/MathGiu/ChaseAndRun. Code for the IBM model is available at https://github.com/jonathan-potts/ChaseAndRun.

## Author Contributions

SB - formal analysis, investigation, methodology, writing-reviewing and editing; VG - formal analysis, investigation, methodology, writing-reviewing & editing; KJP - conceptualisation, formal analysis, investigation, methodology, writing-original draft, writing-reviewing & editing; JRP - formal analysis, investigation, methodology, writing-reviewing and editing

### Funding

KJP acknowledges ‘Miur-Dipartimento di Eccellenza’ funding to the Dipartimento di Scienze, Progetto e Politiche del Territorio (DIST). JRP and VG acknowledge support of Engineering and Physical Sciences Research Council (EPSRC) grant EP/V002988/1 awarded to JRP. SB and VG acknowledge the financial support of GNFM-INdAM through ‘INdAM – GNFM Project’, CUP E53C22001930001.

## Acknowledgements

We thank Alf Gerisch and Nadir Fasola for assistance and advice regarding numerical implementation. KJP and SB are members of INdAM-GNFM.

## References

[1] DJ Sumpter. Collective animal behavior. Princeton University Press, 2010.

[2] E Scarpa and R Mayor. “Collective cell migration in development”. In: Journal of Cell Biology 212.2 (2016), pp. 143–155.

[3] SA Vilchez Mercedes, F Bocci, H Levine, JN Onuchic, MK Jolly, and PK Wong. “Decoding leader cells in collective cancer invasion”. In: Nature Reviews Cancer 21.9 (2021), pp. 592–604.

[4] S Motsch and E Tadmor. “Heterophilious dynamics enhances consensus”. In: SIAM Review 56.4 (2014), pp. 577–621.

[5] AM Berdahl, AB Kao, A Flack, PA Westley, EA Codling, ID Couzin, AI Dell, and D Biro. “Collective animal navigation and migratory culture: from theoretical models to empirical evidence”. In: Philosophical Transactions of the Royal Society B: Biological Sciences 373.1746 (2018), p. 20170009.

[6] KJ Painter, T Hillen, and JR Potts. “Biological modelling with nonlocal advection diffusion equations”. In: Mathematical Models and Methods in Applied Sciences (M3AS) 34 (2023), pp. 57– 107.

[7] JR Potts and MA Lewis. “Spatial memory and taxis-driven pattern formation in model ecosystems”. In: Bulletin of Mathematical Biology 81 (2019), pp. 2725–2747.

[8] CUM Smith. Biology of sensory systems. John Wiley & Sons, 2008.

[9] R Payne and D Webb. “Orientation by means of long range acoustic signaling in baleen whales”. In: Annals of the New York Academy of Sciences 188.1 (1971), pp. 110–141.

[10] YM Yamashita, M Inaba, and M Buszczak. “Specialized intercellular communications via cytonemes and nanotubes”. In: Annual Review of Cell and Developmental Biology 34 (2018), pp. 59–84.

[11] TY Moore and AA Biewener. “Outrun or outmaneuver: predator–prey interactions as a model system for integrating biomechanical studies in a broader ecological and evolutionary context”. In: Integrative and Comparative Biology 55.6 (2015), pp. 1188–1197.

[12] H Yamanaka and S Kondo. “In vitro analysis suggests that difference in cell movement during direct interaction can generate various pigment patterns in vivo”. In: Proceedings of the National Academy of Sciences 111.5 (2014), pp. 1867–1872.

[13] E Theveneau, B Steventon, E Scarpa, S Garcia, X Trepat, A Streit, and R Mayor. “Chase-and-run between adjacent cell populations promotes directional collective migration”. In: Nature Cell Biology 15.7 (2013), pp. 763–772.

[14] GA Vega-Lopez, S Cerrizuela, C Tribulo, and MJ Aybar. “Neurocristopathies: New insights 150 years after the neural crest discovery”. In: Developmental Biology 444 (2018), S110–S143.

[15] NJ Armstrong, KJ Painter, and JA Sherratt. “A continuum approach to modelling cell–cell adhesion”. In: Journal of Theoretical Biology 243.1 (2006), pp. 98–113.

[16] KJ Painter, J Bloomfield, J Sherratt, and A Gerisch. “A nonlocal model for contact attraction and repulsion in heterogeneous cell populations”. In: Bulletin of Mathematical Biology 77 (2015), pp. 1132–1165.

[17] H Murakawa and H Togashi. “Continuous models for cell–cell adhesion”. In: Journal of Theoretical Biology 374 (2015), pp. 1–12.

[18] JA Carrillo, Y Huang, and M Schmidtchen. “Zoology of a nonlocal cross-diffusion model for two species”. In: SIAM Journal on Applied Mathematics 78.2 (2018), pp. 1078–1104.

[19] JA Carrillo, H Murakawa, M Sato, H Togashi, and O Trush. “A population dynamics model of cell-cell adhesion incorporating population pressure and density saturation”. In: Journal of Theoretical Biology 474 (2019), pp. 14–24.

[20] TJ Jewell, AL Krause, PK Maini, and EA Gaffney. “Patterning of nonlocal transport models in biology: the impact of spatial dimension”. In: Mathematical Biosciences 366 (2023), p. 109093.

[21] C Falcú, RE Baker, and JA Carrillo. “A local continuum model of cell-cell adhesion”. In: SIAM Journal on Applied Mathematics (2023), S17–S42.

[22] P Amorim, B Telch, and LM Villada. “A reaction–diffusion predator–prey model with pursuit, evasion, and nonlocal sensing”. In: Mathematical Biosciences and Engineering 16.5 (2019), pp. 5114–5145.

[23] E Ellefsen and N Rodriguez. “On equilibrium solutions to nonlocal mechanistic models in ecology”. In: Journal of Applied Analysis and Computation 11.6 (2021).

[24] V Giunta, T Hillen, M Lewis, and JR Potts. “Local and global existence for nonlocal multispecies advection-diffusion models”. In: SIAM Journal on Applied Dynamical Systems 21.3 (2022), pp. 1686–1708.

[25] V Giunta, T Hillen, MA Lewis, and JR Potts. “Detecting minimum energy states and multistability in nonlocal advection–diffusion models for interacting species”. In: Journal of Mathematical Biology 85.5 (2022), p. 56.

[26] H Wang and Y Salmaniw. “Open problems in PDE models for knowledge-based animal movement via nonlocal perception and cognitive mapping”. In: Journal of Mathematical Biology 86.5 (2023), p. 71.

[27] JR Potts and KJ Painter. “Distinguishing between long-transient and asymptotic states in a biological aggregation model”. In: Bulletin of Mathematical Biology 86.3 (2024), pp. 1–16.

[28] C Carmona-Fontaine, HK Matthews, S Kuriyama, M Moreno, GA Dunn, M Parsons, CD Stern, and R Mayor. “Contact inhibition of locomotion in vivo controls neural crest directional migration”. In: Nature 456.7224 (2008), pp. 957–961.

[29] E Theveneau, L Marchant, S Kuriyama, M Gull, B Moepps, M Parsons, and R Mayor. “Collective chemotaxis requires contact-dependent cell polarity”. In: Developmental Cell 19.1 (2010), pp. 39– 53.

[30] C Carmona-Fontaine, E Theveneau, A Tzekou, M Tada, M Woods, KM Page, M Parsons, JD Lambris, and R Mayor. “Complement fragment C3a controls mutual cell attraction during collective cell migration”. In: Developmental Cell 21.6 (2011), pp. 1026–1037.

[31] A Colombi, M Scianna, KJ Painter, and L Preziosi. “Modelling chase-and-run migration in heterogeneous populations”. In: Journal of Mathematical Biology 80.1 (2020), pp. 423–456.

[32] JW Laundré, L Hernández, and WJ Ripple. “The landscape of fear: ecological implications of being afraid”. In: The Open Ecology Journal 3.1 (2010).

[33] KM Gaynor, JS Brown, AD Middleton, ME Power, and JS Brashares. “Landscapes of fear: spatial patterns of risk perception and response”. In: Trends in Ecology & Evolution 34.4 (2019), pp. 355–368.

[34] HJ Williams, LA Taylor, S Benhamou, AI Bijleveld, TA Clay, S de Grissac, U Demšar, HM English, N Franconi, A Gúmez-Laich, et al. “Optimizing the use of biologgers for movement ecology research”. In: Journal of Animal Ecology 89.1 (2020), pp. 186–206.

[35] G Bastille-Rousseau, JR Potts, JA Schaefer, MA Lewis, EH Ellington, ND Rayl, SP Mahoney, and DL Murray. “Unveiling trade-offs in resource selection of migratory caribou using a mechanistic movement model of availability”. In: Ecography 38.10 (2015), pp. 1049–1059.

[36] TR Ganz, MT DeVivo, AJ Wirsing, SB Bassing, BN Kertson, SL Walker, and LR Prugh. “Cougars, wolves, and humans drive a dynamic landscape of fear for elk”. In: Ecology (2024), e4255.

[37] P Kareiva and G Odell. “Swarms of predators exhibit “preytaxis” if individual predators use area-restricted search”. In: The American Naturalist 130.2 (1987), pp. 233–270.

[38] AC Nisi, JP Suraci, N Ranc, LG Frank, A Oriol-Cotterill, S Ekwanga, TM Williams, and CC Wilmers. “Temporal scale of habitat selection for large carnivores: Balancing energetics, risk and finding prey”. In: Journal of Animal Ecology 91.1 (2022), pp. 182–195.

[39] JM Lee, T Hillen, and MA Lewis. “Pattern formation in prey-taxis systems”. In: Journal of Biological Dynamics 3.6 (2009), pp. 551–573.

[40] S Fagioli and Y Jaafra. “Multiple patterns formation for an aggregation/diffusion predator-prey system”. In: Networks and Heterogeneous Media 16.3 (2021), pp. 377–411.

[41] L Börger, BD Dalziel, and JM Fryxell. “Are there general mechanisms of animal home range behaviour? A review and prospects for future research”. In: Ecology Letters 11.6 (2008), pp. 637– 650.

[42] N Bonnot, N Morellet, H Verheyden, B Cargnelutti, B Lourtet, F Klein, and AM Hewison. “Habitat use under predation risk: hunting, roads and human dwellings influence the spatial behaviour of roe deer”. In: European Journal of Wildlife Research 59 (2013), pp. 185–193.

[43] J Krause, D Hoare, S Krause, C Hemelrijk, and D Rubenstein. “Leadership in fish shoals”. In: Fish and Fisheries 1.1 (2000), pp. 82–89.

[44] SA Rands, G Cowlishaw, RA Pettifor, JM Rowcliffe, and RA Johnstone. “Spontaneous emergence of leaders and followers in foraging pairs”. In: Nature 423.6938 (2003), pp. 432–434.

[45] R McLennan, LJ Schumacher, JA Morrison, JM Teddy, D. Ridenour, AC Box, CL Semerad, H Li, W McDowell, D Kay, et al. “Neural crest migration is driven by a few trailblazer cells with a unique molecular signature narrowly confined to the invasive front”. In: Development 142.11 (2015), pp. 2014–2025.

[46] S Nakayama, JL Harcourt, RA Johnstone, and A Manica. “Who directs group movement? Leader effort versus follower preference in stickleback fish of different personality”. In: Biology Letters 12.5 (2016), p. 20160207.

[47] E Theveneau and C Linker. “Leaders in collective migration: are front cells really endowed with a particular set of skills?” In: F1000Research 6 (2017).

[48] E Scarpa, A Roycroft, E Theveneau, E Terriac, M Piel, and R Mayor. “A novel method to study contact inhibition of locomotion using micropatterned substrates”. In: Biology Open 2.9 (2013), pp. 901–906.

[49] A Szabú, M Melchionda, G Nastasi, ML Woods, S Campo, R Perris, and R Mayor. “In vivo confinement promotes collective migration of neural crest cells”. In: Journal of Cell Biology 213.5 (2016), pp. 543–555.

[50] R McLennan, R Giniunaite, K Hildebrand, JM Teddy, JC Kasemeier-Kulesa, L Bolanos, RE Baker, PK Maini, and PM Kulesa. “Colec12 and Trail signaling confine cranial neural crest cell trajectories and promote collective cell migration”. In: Developmental Dynamics 252.5 (2023), pp. 629–646.

[51] V Giunta, T Hillen, MA Lewis, and JR Potts. “Weakly nonlinear analysis of a two-species nonlocal advection–diffusion system”. In: Nonlinear Analysis: Real World Applications 78 (2024), p. 104086.

[52] A Gerisch and KJ Painter. “Mathematical modelling of cell adhesion and its applications to developmental biology and cancer invasion”. In: vol. 2. CRC Press Boca Raton, 2010, pp. 319– 350.

[53] TYC Tsai, RM Garner, and SG Megason. “Adhesion-based self-organization in tissue patterning”. In: Annual Review of Cell and Developmental Biology 38 (2022), pp. 349–374.

