## Supplementary Information for "Variations in nonlocal interaction range lead to emergent chase-and-run in heterogeneous populations"

[1] *Dipartimento Interateneo di Scienze, Progetto e Politiche del Territorio (DIST), Politecnico di Torino, Viale Pier Andrea Mattioli 39, 10125, Turin, Italy*
[2] *Department of Mathematics, Swansea University, Computational Foundry, Bay Campus, Swansea SA1 8EN , UK*
[3] *School of Mathematics and Statistics, Hounsfield Road, University of Sheffield, UK, S3 7RH*
[4] *Department of Mathematical Sciences "G. L. Lagrange", Politecnico di Torino, Corso Duca degli Abruzzi 24, 10129 Torino, Italy*


### A Individual based model

#### A.1 Model description

Here we describe the stochastic individual based model (IBM); as noted in the main text, the IBM is constructed in a manner that permits formal relation to the continuous model (Equations (1) in main manuscript) in an appropriate limit, demonstrated below. The IBM takes the form of a position jump random walk on a lattice, where we work on either a 1D line lattice of length $L$ or a 2D square lattice of side $L$, with periodic boundary conditions. The lattice spacing is denoted $l$ and defined so that $L/l$ is an integer. We denote by $M(x,t)$ (resp. $N(x,t)$) the number of individuals from population $C$ (resp. $R$) at lattice point $x$ at time $t$. For the 1D model, we define

$$S^1_M(x,t) = \min\Bigg\{1, \max\Bigg\{-1, \phi\left(\frac{M(x)+N(x)}{l}\right)\left(\frac{\alpha_{CC}}{4r_{CC}}\sum_{y=-r_{CC}}^{r_{CC}} M(x+yl,t)\frac{y}{|y|}\right.$$
$$\left. + \frac{\alpha_{CR}}{4r_{CR}}\sum_{y=-r_{CR}}^{r_{CR}} N(x+yl,t)\frac{y}{|y|}\right)\Bigg\}\Bigg\}, \quad (1)$$

$$S^1_N(x,t) = \min\Bigg\{1, \max\Bigg\{-1, \phi\left(\frac{M(x)+N(x)}{l}\right)\left(\frac{\alpha_{RR}}{4r_{RR}}\sum_{y=-r_{RR}}^{r_{RR}} N(x+yl,t)\frac{y}{|y|}\right.$$
$$\left. + \frac{\alpha_{RC}}{4r_{RC}}\sum_{y=-r_{RC}}^{r_{RC}} M(x+yl,t)\frac{y}{|y|}\right)\Bigg\}\Bigg\}, \quad (2)$$

where $r_{CC}, r_{CR}, r_{RC}, r_{RR} < L/l$ are integers. In 2D, we define

$$S^2_{M1}(x,t) = \min\Bigg\{1, \max\Bigg\{-1, \phi\left(\frac{M(x)+N(x)}{l^2}\right)\left(\frac{\alpha_{CC}}{2\pi l r^2_{CC}}\sum_{\mathbf{y}\in B^2_{r_{CC}}} M(\mathbf{x}+\mathbf{y}l,t)\cos(\theta_{\mathbf{y}})\right.$$
$$\left. + \frac{\alpha_{CR}}{2\pi l r^2_{CR}}\sum_{\mathbf{y}\in B^2_{r_{CR}}} N(\mathbf{x}+\mathbf{y}l,t)\cos(\theta_{\mathbf{y}})\right)\Bigg\}\Bigg\}, \quad (3)$$

$$S^2_{M2}(x,t) = \min\Bigg\{1, \max\Bigg\{-1, \phi\left(\frac{M(x)+N(x)}{l^2}\right)\left(\frac{\alpha_{CC}}{2\pi l r^2_{CC}}\sum_{\mathbf{y}\in B^2_{r_{CC}}} M(\mathbf{x}+\mathbf{y}l,t)\sin(\theta_{\mathbf{y}})\right.$$
$$\left. + \frac{\alpha_{CR}}{2\pi l r^2_{CR}}\sum_{\mathbf{y}\in B^2_{r_{CR}}} N(\mathbf{x}+\mathbf{y}l,t)\sin(\theta_{\mathbf{y}})\right)\Bigg\}\Bigg\}, \quad (4)$$

where $\theta_{\mathbf{y}}$ is the direction of the vector $\mathbf{y}$. Functions $S^2_{N1}(x,t)$ and $S^2_{N2}(x,t)$ are defined analogously.

For the 1D random walk we assume that, in a time-step of length $\tau$, there is some probability that an individual at $x$ moves to one of the two adjacent lattice sites at $x \pm l$. For a member of population $C$, these probabilities are given by

$$p^1_{M\tau}(x \pm l|x) = \frac{1 \pm S^1_M(x,t)}{2}, \quad (5)$$

with an analogous expression for the probabilities of movement for members of population $R$.

In the 2D random walk, movements can take place to one of the four adjacent lattice sites, i.e. from position $\mathbf{x}$ to $\mathbf{x} \pm (l, 0)$ or $\mathbf{x} \pm (0, l)$. For a member of population $C$, these probabilities are given by

$$p^2_{M\tau}(\mathbf{x} \pm (l,0)|\mathbf{x}) = \frac{1 \pm S^2_{M1}(\mathbf{x},t)}{4}, \tag{6}$$

$$p^2_{M\tau}(\mathbf{x} \pm (0,l)|\mathbf{x}) = \frac{1 \pm S^2_{M2}(\mathbf{x},t)}{4}, \tag{7}$$

with analogous expressions for the probabilities of movement for members of population $R$. To perform simulations, the IBMs were coded in C and the code can be found at `https://github.com/jonathan-potts/ChaseAndRun`.

#### A.2 From IBM to PDE

To relate the IBMs described in Equations (1)-(7) to the PDEs of Equations (1) in the main manuscript, we first take expectations and assume that covariances are negligible (i.e. a mean field approximation). With this assumption, the expected number of individuals, $M_E(x,t)$, from population $C$ at lattice site $x$ at time $t$ obeys the following iterative equation in 1D

$$\begin{aligned} M_E(x,t+\tau) &= M_E(x-l,t)p^1_{C\tau}(x|x-l) + M_E(x+l,t)p^1_{C\tau}(x|x+l) \\ &= \frac{1}{2}[M_E(x-l,t)(1+S^1_M(x-l,t)) + M_E(x+l,t)(1-S^1_M(x+l,t))], \end{aligned} \tag{8}$$

and the equation for $N_E(x,t)$ (the expected number of individuals from population $R$ at lattice site $x$ at time $t$) is analogous. In 2D, the change in $M_E(x,t)$ over time is given by

$$\begin{aligned} M_E(\mathbf{x},t+\tau) =& \frac{1}{4}[M_E(\mathbf{x}-(l,0),t)(1+S^2_{M1}(\mathbf{x}-(l,0),t)) + M_E(\mathbf{x}+(l,0),t)(1-S^2_{M1}(\mathbf{x}+(l,0),t)) \\ &+ M_E(\mathbf{x}-(0,l),t)(1+S^2_{M2}(\mathbf{x}-(0,l),t)) + M_E(\mathbf{x}+(0,l),t)(1-S^2_{M2}(\mathbf{x}+(0,l),t))]. \end{aligned} \tag{9}$$

For the 1D case, we rearrange Equation (8) to give

$$\begin{aligned} \frac{M_E(x,t+\tau) - M_E(x,t)}{\tau} = \frac{l^2}{2\tau}\Bigg[&\frac{M_E(x+l,t) - 2M_E(x,t) + M_E(x-l,t)}{l^2} \\ &- \frac{2}{l}\frac{S^1_M(x+l,t)M_E(x+l,t) - S^1_M(x-l,t)M_E(x-l,t)}{2l}\Bigg]. \end{aligned} \tag{10}$$

In 2D, a similar rearrangement gives

$$\begin{aligned} &\frac{M_E(\mathbf{x},t+\tau) - M_E(\mathbf{x},t)}{\tau} = \\ &\frac{l^2}{2\tau}\Bigg[\frac{M_E(\mathbf{x}+(l,0),t) + M_E(\mathbf{x}+(0,l),t) - 4M_E(\mathbf{x},t) + M_E(\mathbf{x}-(l,0),t) + M_E(\mathbf{x}-(0,l),t)}{l^2} \\ &- \frac{2}{l}\frac{S^2_{M1}(\mathbf{x}+(l,0),t)M_E(\mathbf{x}+(l,0),t) - S^2_{M1}(\mathbf{x}-(l,0),t)M_E(\mathbf{x}-(l,0),t)}{2l} \\ &- \frac{2}{l}\frac{S^2_{M2}(\mathbf{x}+(0,l),t)M_E(\mathbf{x}+(0,l),t) - S^2_{M2}(\mathbf{x}-(0,l),t)M_E(\mathbf{x}-(0,l),t)}{2l}\Bigg]. \end{aligned} \tag{11}$$

We then take the limit as $l, \tau \to 0$ such that $d_n = l^2/(2n\tau)$ is kept constant (i.e. the diffusion limit). We also send $r_{ij} \to \infty$ keeping $\xi_{ij} = r_{ij}l$ constant, for each $i, j \in \{C, R\}$ and write $C(\mathbf{x},t)$ (resp. $R(\mathbf{x},t)$) for the limit of $M_E(\mathbf{x}_l,t)/l^n$ (resp. $N_E(\mathbf{x}_l,t)/l^n$) where $\mathbf{x}_l$ is the closest lattice site to the point $\mathbf{x}$ for any given $l$ (using bold letters for both 1D and 2D here). A direct calculation of this limit leads to

$$\begin{aligned} \frac{\partial C}{\partial t} &= d_n\nabla\cdot\left[\nabla C - \frac{C\alpha_{CC}}{V_n(\xi_{CC})}\phi(C+R)\int_{\mathcal{B}^n_{\xi_{CC}}} C(\mathbf{x}+\mathbf{y},t)e_\mathbf{y}\,d\mathbf{y} - \frac{C\alpha_{CR}}{V_n(\xi_{CR})}\phi(C+R)\int_{\mathcal{B}^n_{\xi_{CR}}} R(\mathbf{x}+\mathbf{y},t)e_\mathbf{y}\,d\mathbf{y}\right], \\ \frac{\partial R}{\partial t} &= d_n\nabla\cdot\left[\nabla R - \frac{R\alpha_{RC}}{V_n(\xi_{RC})}\phi(C+R)\int_{\mathcal{B}^n_{\xi_{RC}}} C(\mathbf{x}+\mathbf{y},t)e_\mathbf{y}\,d\mathbf{y} - \frac{R\alpha_{RR}}{V_n(\xi_{RR})}\phi(C+R)\int_{\mathcal{B}^n_{\xi_{RR}}} R(\mathbf{x}+\mathbf{y},t)e_\mathbf{y}\,d\mathbf{y}\right]. \end{aligned} \tag{12}$$

In the case $d_n = 1$, this is the same as Equations (1) in the main manuscript for $D_C = D_R = 1$. The case $D_C = D_R = d_n = 1$ is the only case we are interested in for our numerical analysis. However, note that one could set $d \neq 0$ and rescale the $\alpha'_{ij}s$ to return Equations (1) of the manin manuscript for any non-zero values of $D_C$ and $D_R$, if required. We leave this general case as an exercise for any interested reader.

Finally, when comparing between IBM and PDE formulation, it is possible rescale by any constant $A$, by setting $C \mapsto C/A$, $R \mapsto R/A$, and $\alpha_{ij} \mapsto A\alpha_{ij}$, for $i, j \in \{C, R\}$. This can be valuable, as the total mass in the IBM version is by definition the number of individuals. Therefore this rescaling allows comparisons can be made between a single set of PDEs, where the total mass of $A$ and $B$ is fixed across all the PDE analysis, and corresponding IBMs with different numbers of individuals.

### B Linear Stability Analysis

Following the standard approach (e.g. [1]), we consider small heterogeneous perturbations of the spatially uniform steady state $(C_s, R_s)$, i.e. $C(\mathbf{x}, t) = C_s + \tilde{C}(\mathbf{x}, t)$ and $R(\mathbf{x}, t) = R_s + \tilde{R}(\mathbf{x}, t)$, with $|\tilde{C}(\mathbf{x}, t)|, |\tilde{R}(\mathbf{x}, t)| \ll 1$. Substituting into Equations (1) of the main manuscript, linearising about $(C_s, R_s)$, and looking for solutions of the form $\tilde{C}, \tilde{R} \propto e^{i\mathbf{k}\cdot\mathbf{x}+\lambda t}$ (where $\mathbf{k}$ denotes the wave-vector and $\lambda$ is the growth rate) leads to the dispersion relation

$$\lambda^2 + \mathcal{C}(k)\lambda + \mathcal{D}(k) = 0, \tag{13}$$

where $\mathcal{C}(k)$ and $\mathcal{D}(k)$ are given by

$$\mathcal{C}(k) = k^2(D_C + D_R) - k\,(\Lambda_{CC} + \Lambda_{RR}) \tag{14a}$$
$$\mathcal{D}(k) = D_C\,D_R\,k^4 - k^3\,(D_R\,\Lambda_{CC} + D_C\,\Lambda_{RR}) + k^2\,(\Lambda_{CC}\,\Lambda_{RR} - \Lambda_{CR}\Lambda_{RC})\,. \tag{14b}$$

Each $\Lambda_{uv}$ function represents the nonlocal contribution for each interaction and is evaluated as proposed in [2], where the authors extend pattern formation analysis to higher spatial dimension for models of form (**??**). They are given by:

$$\Lambda_{UV} = U_s\,\frac{2\pi^{\frac{n}{2}}}{\Gamma(\frac{n}{2})}\,\frac{\alpha_{UV}}{V_n(\xi_{UV})}\,\phi(U_s + V_s)\int_0^{\xi_{UV}} y^{n-1}\,j_1^{(n)}(ky)\,dy, \tag{15}$$

where $k = |\mathbf{k}|$, $y = |\mathbf{y}|$ and $U, V \in \{C, R\}$ with corresponding homogeneous steady state $U_s, V_s \in \{C_s, R_s\}$. In the expression above, $j_1^{(n)}(\mathbf{x})$ denote the first order $n^{\text{th}}$ dimensional hyperspherical Bessel functions. In particular, $j_1^{(1)}(x) = \sin(x)$ and $j_1^{(2)}(x) = J_1(kx)$, where $J_1$ denotes the first order Bessel function of the first kind.

In 1D ($n = 1$), after a few rearrangements, the nonlocal terms in Eq. (15) reduce to

$$\Lambda_{UV} = U_s\alpha_{UV}\phi(U + V)\Gamma_{UV}, \quad \text{where } \Gamma_{UV} = \frac{1 - \cos(\xi_{UV}\,k)}{\xi_{UV}\,k}, \tag{16}$$

for $U, V \in \{C, R\}$, $U_s \in \{C_s, R_s\}$.

In 2D ($n = 2$), we instead have

$$\Lambda_{UV} = 2U_s\frac{\alpha_{UV}}{\xi_{UV}^2}\phi(U_s + V_s)\left(-\frac{\xi_{UV}}{k}J_0(k\xi_{UV}) + \frac{1}{k^2}\int_0^{k\xi_{UV}} J_0(p)dp\right), \tag{17}$$

where $J_0$ is the zero order Bessel function of the first kind.

To assess the potential for pattern formation, we investigate when the uniform steady state is stable to spatially homogeneous perturbations (i.e. $\Re(\lambda_+(0)) \leq 0$, where $\lambda_+ = 0.5(-\mathcal{C} + \sqrt{\mathcal{C}^2 - 4\mathcal{D}})$) and unstable to spatially inhomogeneous perturbations (i.e. $\Re(\lambda_+(k)) > 0$, for some $k > 0$ – we refer to any such $k$ as unstable wavenumbers). The former condition is always satisfied. The latter holds if and only if $\mathcal{C}(k) < 0$ or $\mathcal{D}(k) < 0$, for some positive $k$. The tractability of the 1D dispersion relation allows for the derivation of some general insights. Specifically, in 1D (using the simplified form $\phi \equiv 1$ for consistency with our numerics) the condition on $\mathcal{C}$ gives

$$C_s\,\alpha_{CC}\,\xi_{CC} + R_s\,\alpha_{RR}\,\xi_{RR} > 2\,(D_C + D_R), \tag{18}$$

which shows that an instability is possible when the positive homotypic interactions dominate over diffusion: i.e. at least one of the populations has a sufficiently strong self-attraction.

Moreover, for instances in which pattern formation is predicted, we distinguish between *Turing instabilities* (i.e. $\Im(\lambda_+(k)) = 0$ for unstable wavenumbers) and *Turing-wave instabilities* (when $\Im(\lambda_+(k)) \neq 0$ for at least one of the unstable wavenumbers); in the latter, solutions are expected to oscillate in space and time as they diverge from the uniform steady state. Turing-wave patterns require $\mathcal{C}^2(k) - 4\mathcal{D}(k) < 0$ and, for the 1D model with $\phi \equiv 1$ and $D_C = D_R = 1$, this leads to

$$(C_s\,\alpha_{CC}\,\xi_{AA} - R_s\,\alpha_{RR}\,\xi_{RR})^2 + 4C_s\,R_s\,\alpha_{CR}\,\xi_{CR}\,\alpha_{RC}\,\xi_{RC} < 0, \tag{19}$$

Thus, in 1D, Turing-wave patterns may be possible under a sufficiently strong chase-and-run interaction to emerge.

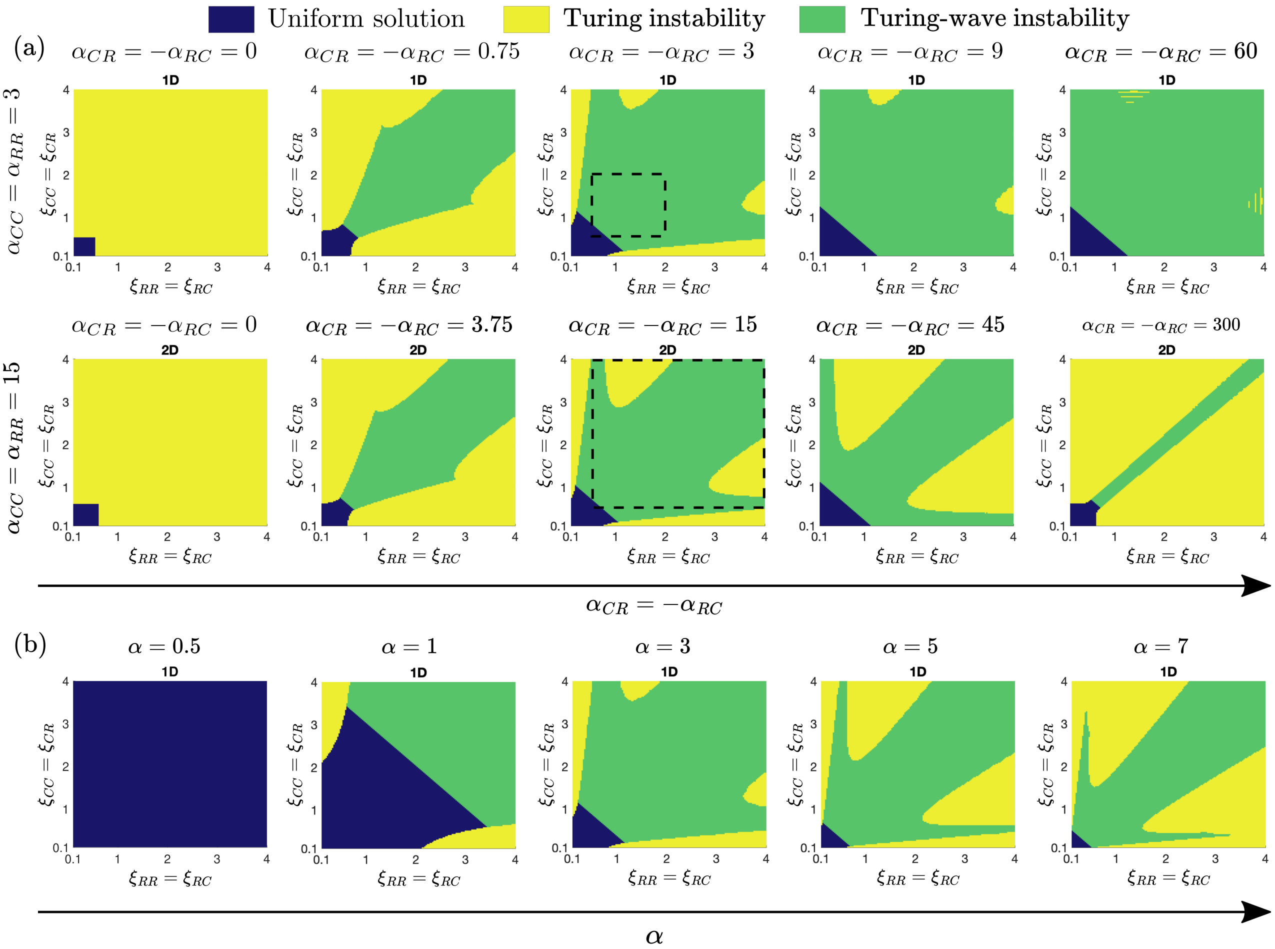


Figure S1: Parameter spaces for pattern formation predicted by the LSA in 1D and 2D, for increasing values of (a) the chase-and-run interactions and (b) all interactions ranges. We classify the resulting solutions as: uniform solution (blue), stationary pattern (yellow), dynamic pattern (green). The dotted boxes highlight the values considered in the numerical studies presented in the main text, compare respectively with Figure 3(a) (1D) and Figure 4(a) (2D). We adopt $\phi(C, R) = 1$ in 1D, $\phi(C, R) = 1/(1 + C + R)$ in 2D. Other model parameters are set as $D_C = D_R = 1$, $C_s = R_s = 1$.

In Figure S1 we plot the predicted parameter space for patterning in both 1D (using $\phi \equiv 1$) and 2D (using $\phi \equiv 1/(1 + C + R)$) across $(\xi_R, \xi_C)$ space. Note that the middle panels in Figure S1(a) in the top and bottom rows correspond to the simulation study settings of Figure 3(a) and Figure 4(a), respectively. In both instances, the regions of predicted patterning match with the results of the numerical simulations. Moreover, the growth rate – by which we mean $\max\{\Re(\lambda_+(k))\}$, an indicator of how quickly solutions are expected to diverge from the steady state – is found to increase if we move away from the stability region (path (i)-(ii)-(iii)) of Figure 3(a) (result not shown). From left to right we show the impact of an increasingly strong chase-and-run interaction which, for a moderate interaction leads to a broad transition from Turing to Turing-wave type instabilities. A more dominant chase-and-run, though, has a diverging impact in 1D and 2D, where in the former we observe that in 1D patterns are near ubiquitously predicted to be of Turing-wave like, while in 2D they are of Turing-like. This is consistent with the significant differences between 1D and 2D noted previously in [2]. Finally we note that if we simultaneously increase all interaction strengths, any regions of uniform solutions are found to reduce and instabilities are predicted to form under smaller interaction ranges, see Figure S1(b).

### C Numerical methods

For the 1D simulations we adopt a Method of Lines approach. Specifically, we discretise the domain $[0, L]$ into a regular lattice of spacing $\Delta x$ and solve the resultant system of time-dependent ODEs. Note that the discretisation of the nonlocal terms exploits a Fast Fourier Transform technique to efficiently calculate the integral: full details of the numerical method itself are provided in [3]. For all 1D simulations we have set $\Delta x = 0.05$. The code used to solve Equations (1) of the main manuscript in 1D is available at `https://github.com/kjpainter/ChaseAndRun`. For the 2D simulations we use the spectral numerical scheme described in [4]. We discretise the spatial domain

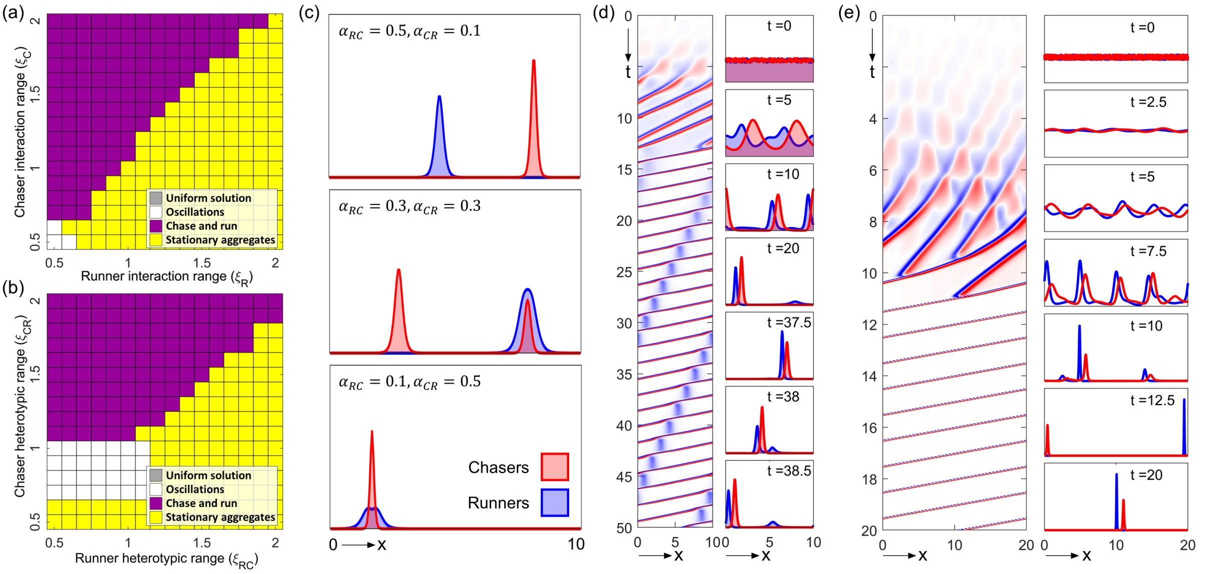


Figure S2: (a) Pattern selection across interaction range space. The form of pattern is classified following a simulation at each $(\xi_C, \xi_R)$ pair as: uniform solution (gray), spatiotemporal oscillations (white), population chase-and-run (magenta), stationary aggregates (yellow). Here, $\alpha_{CC} = \alpha_{CR} = -\alpha_{RC} = \alpha_{RR} = 5$. (b) Pattern selection across heterotypic interaction range space $(\xi_{RC}, \xi_{CR})$. We set $\xi_{CC} = \xi_{RR} = 1$ and $\alpha_{CC} = \alpha_{CR} = -\alpha_{RC} = \alpha_{RR} = 3$. (c) Different forms of stationary aggregate when $1 = \xi_C < \xi_R = 1.5$ and $\alpha_{CC} = \alpha_{RR} = 3$: (top) completely segregated, $\alpha_{CR} = 1$ $\alpha_{RC} = -5$; (middle) mixed/segregated, $\alpha_{CR} = 3$ $\alpha_{RC} = -3$; (bottom) completely mixed, $\alpha_{CR} = 5$ $\alpha_{RC} = -1$. (d) Chase-and-run, where a group of runners is periodically dropped. $\xi_C = 1.5$, $\xi_R = 1$, $\alpha_{CC} = \alpha_{CR}, \alpha_{RR} = 3$, $\alpha_{RC} = -1.5$. (e) Chase-and-run on a larger domain $\xi_C = 1.5, \xi_R = 1$, $\alpha_{CC} = \alpha_{CR} = \alpha_{RR} = 3, \alpha_{RC} = -3$. For all simulations in this figure, $D_C = D_R = R_S = C_S = 1$ and initial conditions are dispersed on a domain of length $L = 10$ for panels (a) - (d), and $L = 20$ for panel (e).

$[0, L] \times [0, L]$ by defining the grid points $(x_i, y_j)$, where $x_i = i\Delta x$, $y_j = j\Delta y$, and $i, j \in \{0, 1, \ldots, 2^n - 1\}$, with $n = 7$ for $L = 10$, and $n = 8$ for $L = 20$. The code used to solve Equations (1) of the main manuscript in 2D is available at `https:https://github.com/MathGiu/ChaseAndRun`.

### D Additional figures

### References


[1] JD Murray. *Mathematical biology II: spatial models and biomedical applications.* Vol. 18. Springer, 2003.

[2] TJ Jewell, AL Krause, PK Maini, and EA Gaffney. “Patterning of nonlocal transport models in biology: the impact of spatial dimension”. In: *Mathematical Biosciences* 366 (2023), p. 109093.

[3] A Gerisch. “On the approximation and efficient evaluation of integral terms in PDE models of cell adhesion”. In: *IMA journal of numerical analysis* 30.1 (2010), pp. 173–194.

[4] V Giunta, T Hillen, M Lewis, and JR Potts. “Local and global existence for nonlocal multispecies advection-diffusion models”. In: *SIAM Journal on Applied Dynamical Systems* 21.3 (2022), pp. 1686–1708.

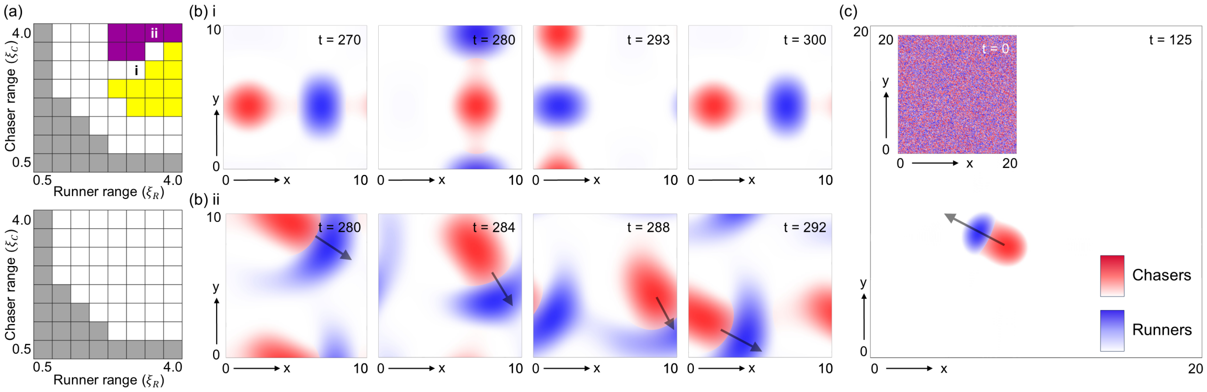


Figure S3: (a) Pattern selection across $(\xi_R, \xi_C)$ space for 2D simulations. The form of pattern is classified following a simulation at each $(\xi_R, \xi_C)$ pair as: uniform solution (gray), spatiotemporal oscillations (white), population chase-and-run (magenta), stationary aggregates (yellow). For the top panel we consider uniformly weak interaction strengths ($\alpha_{CC} = \alpha_{CR} = -\alpha_{RC} = \alpha_{RR} = 6$), while for the bottom panel homotypic interaction strengths are weak ($\alpha_{CC} = \alpha_{RR} = 6$) but heterotypic interaction strengths are strong ($\alpha_{CR} = -\alpha_{RC} = 20$). (b) Snapshots for the simulations marked in (a, top panel), showing (i) periodic oscillations; (ii) chase-and-run. (c) Simulation of the 2D model on a larger domain, showing that chase-and-run still forms robustly – the configuration shown at $t = 125$ persistently moves in the direction of the arrow with constant shape and speed. Note that the interaction strength and ranges for this simulation are as in Figure 4. For all simulations in this figure $D_C = D_R = R_s = C_s = 1$ and initial conditions are dispersed.
